# Mapping the micro-proteome of the nuclear lamina and lamin associated domains

**DOI:** 10.1101/828210

**Authors:** Jevon A. Cutler, Xianrong Wong, Victoria E. Hoskins, Molly Gordon, Anil K. Madugundu, Akhilesh Pandey, Karen L. Reddy

## Abstract

The nuclear lamina is a proteinaceous network of filaments that provide both structural and gene regulatory functions by tethering proteins and large domains of DNA, so-called lamin associated domains (LADs), to the periphery of the nucleus. LADs are a large fraction of the mammalian genome that are repressed, in part, by their association to the nuclear periphery. The genesis and maintenance of LADs is poorly understood as are the proteins that participate in these functions. In an effort to identify proteins that reside at the nuclear periphery and potentially interact with LADs, we have taken a two-pronged approach. First, we have undertaken an interactome analysis of the inner nuclear membrane bound LAP2β to further characterize the nuclear lamina proteome. To accomplish this, we have leveraged the BioID system, which previously has been successfully used to characterize the nuclear lamina proteome. Second, we have established a system to identify proteins that bind to LADs by developing a chromatin directed BioID system. We combined the BioID system with the m6A-tracer system which binds to LADs in live cells to identify both LAD proximal and nuclear lamina proteins. In combining these datasets, we have further characterized the protein network at the nuclear lamina, identified putative LAD proximal proteins and found several proteins that appear to interface with both micro-proteomes. Importantly, several proteins essential for LAD function, including heterochromatin regulating proteins related to H3K9 methylation, were identified in this study.

## Introduction

DNA and proteins are highly organized within the eukaryotic cell nucleus. Sequestration of proteins into nuclear sub-domains and the higher order organization of chromatin itself have been implicated in the regulation of the genome [1–6]. Association of chromatin with the nuclear periphery, in particular, has been implicated in gene regulation, specifically correlating with repression of developmentally regulated loci [4,7,8]. DamID (DNA Adenine Methyltransferase Identification), a genome-wide technique to identify nuclear lamina-proximal chromatin, has allowed the identification of Lamina Associated Domains (LADs)[9]. These 100 kilobase (kb) to megabase (Mb) sized domains are enriched for genes that are transcriptionally silent and enriched in histone modifications indicative of facultative heterochromatin, such as histone H3 lysine 9 di/tri-methylation (H3K9me2/3) and histone H3 lysine 27 trimethylation (H3K27me3)[10–13]. Moreover, recent studies have highlighted that both H3K9me2/3 and H3K27me3 are involved in LAD organization [12–16]. LADs represent a large fraction of the genome (30-50%, depending upon the cell type) and are highly correlated with the so-called heterochromatic “B-compartment”, as identified by chromatin conformation capture assays (HiC) [13,17,18]. Given their importance to cellular function and identity, it is important to understand how these large domains of heterochromatin are regulated, maintained, and formed in order to understand global genome regulation and organization. An important element of understanding LAD organization and function is identifying which proteins are present at LADs and the nuclear lamina.

In addition to H3K27me3 and H3K9me2/3, A-type lamins and the inner nuclear membrane (INM) proteins lamina-associated polypeptide beta (LAP2β) and Lamin B Receptor (LBR) have been implicated in organizing LADs [12,17–20]. The nuclear lamina is a proteinaceous network of type V intermediate filaments comprising A and B type lamins. These coiled-coil domain proteins provide a structural scaffold at the INM, with the A-type lamins being shown to mediate LAD organization [12,17–19]. Longstanding efforts have been undertaken to map and characterize the local proteome of the nuclear lamina of the INM using inner nuclear membrane preparations, co-immunoprecipitation and, more recently, BioID (Biotinylation Identification), a method for detecting proximal protein interactions in living cells [21–25]. However, these efforts have exclusively focused on protein members of the INM/lamina as baits and have not measured the protein landscape of LADs themselves or the intersection of these proteome environments. To better understand these proximal protein compartments, we have leveraged the BioID system to study the “micro-proteome” of both LADs and the nuclear lamina using a multi-pronged approach. First, we have generated a chimeric protein comprising a modified promiscuous biotin ligase, BirA*, fused to the nucleoplasmic N-terminus of LAP2β to analyze the interactome of this important protein of the INM/lamina (Figure 1A)[22,23]. Second, we have taken a novel approach combining the specificity of a DamID-based system to label LADs in live cells, the m6A-tracer system, and coupled this with BioID to characterize proteins proximal to LADs (Figure 1B, C) [16]. Finally, we integrate these proteomic data with previous findings on lamin protein interactions to generate a comprehensive mapping of the LAD/lamina proteome interface.

**Figure 1:**
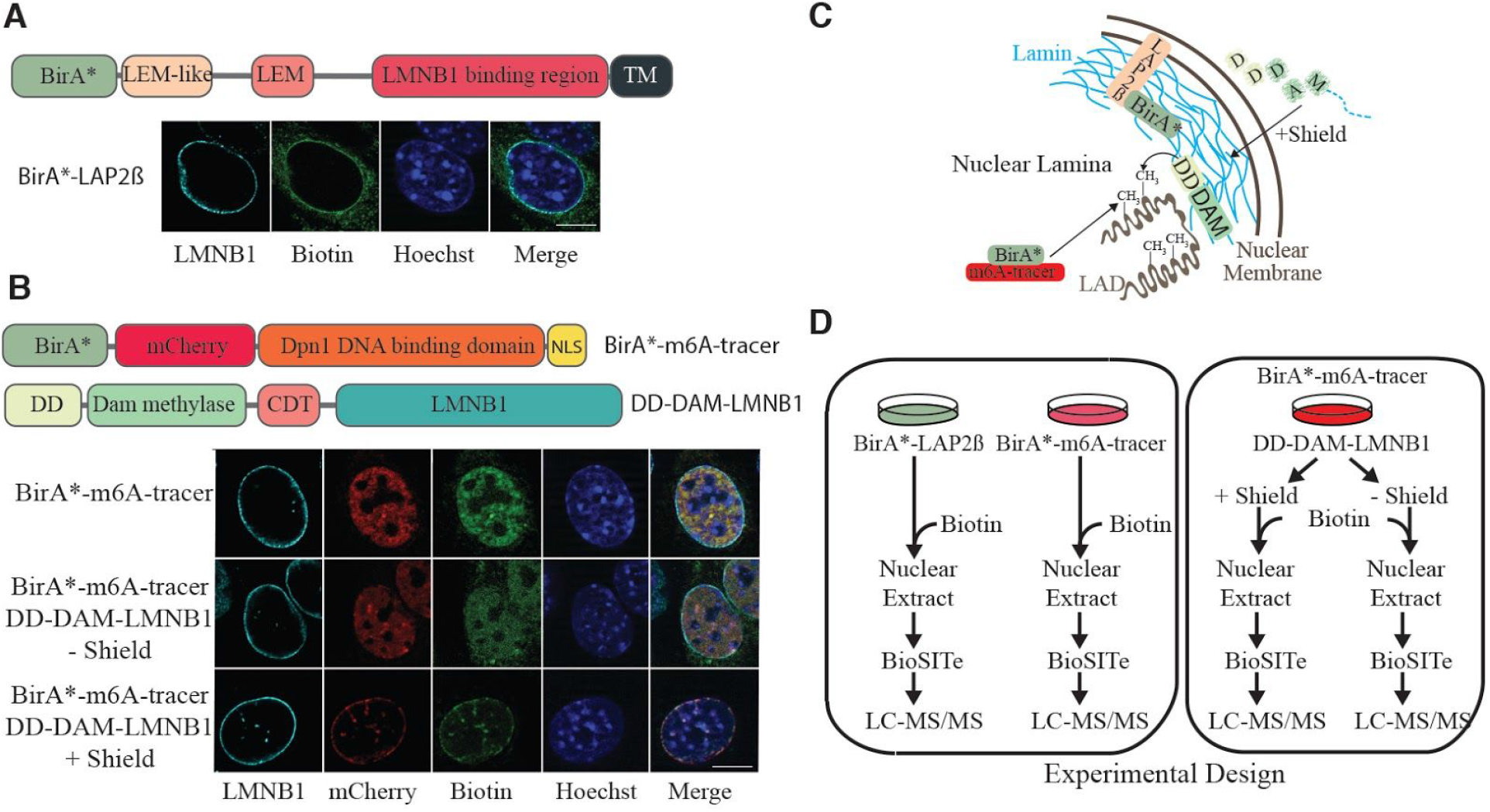
Strategy for investigating the nuclear periphery microproteomes. (A) Domain structures of BirA*-LAP2β and representative images showing the biotinylated interactome of Lap2β (green) at the nuclear periphery. (B) Design of the m6a tracer system for probing the LAD microproteome (top) and representative images showing the expected localization of the BirA*-m6a-tracer (red) and its biotinylated proteins (green) in the absence of DD-DamLmnB1 (top row), in the absence (middle row) and presence (bottom row) of the shield ligand. (C) Pictorial representation of BirA* localization within the nucleus. (D) Experimental workflow for mass spectrometry based BioID LAD microproteome analysis. Cells expressing BirA*-LAP2β, BirA*-m6A-tracer alone, BirA*-m6A-tracer/DD-Dam-LMNB1 plus/minus shield ligand were cultured with exogenous biotin then nuclei were extracted and Biotinylation Site Identification Technology (BioSITe) coupled to liquid chromatography/tandem mass spectrometry analysis was used to identify biotinylated proteins.

While a previous study used CRISPR-directed proximity-labeling approach to identify proteomes associated with repetitive sequences, this study represents, for the first time to our knowledge, a chromatin-directed-BioID strategy that is independent of identifying specific repetitive DNA sequences and targets a specific nuclear compartment [26]. Herein, we identify the proteome of a chromatin compartment (LADs/B-compartment) by leveraging two proximity labeling techniques: BioID and DamID. In addition, by integrating published datasets, our new LAP2β interactome data, and our LAD directed proteome data, we have identified different interaction zones at the nuclear periphery, thus mapping the differential and overlapping microproteomes of the peripheral nuclear compartment. In zone 1, we identified proteins that appear to be restricted to the INM/lamina that do not interact with LADs. In zone 2 we identified proteins that interact with both LADs and the INM/lamina and these may comprise the “middlemen” required to organize the LADs to the nuclear lamina. Finally, in zone 3 we identify proteins that are restricted to LADs, many of which are involved in regulating histone H3 lysine 9 methylation (H3K9me2/3) and cell cycle regulation.

## Results and Discussion

### Establishment of BioID system to map the local proteomes at the nuclear periphery

The BioID system was initially developed using the lamin A protein as the bait and, for the first time, a robust interactome of this insoluble protein was obtained [23]. The BioID system relies on an engineered biotin ligase, BirA* which, when expressed in cells, has a small biotinylation proximity radius, thus labeling lysines on both proximal and directly interacting proteins. BioID, among other methods, has been used to analyze the local proteome of the INM using lamin A/C proteoforms, lamin B1 (LMNB1), SUN domain-containing protein 2 (SUN2), and nuclear pore complex (NPC) members as baits [21,23,27,28]. To expand on these efforts and to focus on an INM protein implicated in LAD organization, we chose to identify the interactome of the lamina-associated polypeptide beta (LAP2β) in a BioID study. The LAP2β protein results from alternative splicing of the gene thymopoietin (*TMPO*) and was initially observed in nuclear envelope isolations and shown to bind lamin proteins [29]. LAP2β is an integral INM protein that is thought to link the nuclear membrane to chromatin and also to regulate transcription factor functions [30]. LAP2β is a member of the LEM (Lap2-Emerin-Man1) domain family of proteins and contains a LEM domain as well as a LEM-like domain, both of which have been thought to mediate interactions between protein/DNA complexes with the lamin binding region. To leverage the BioID system for analysis of the LAP2β proximal interactome we have tagged the nucleoplasmic N-terminus with BirA* (Figure 1A).

BioID is closely related to the DamID technique in that both techniques rely on in-cell labeling of proximal molecules [9]. In DamID, instead of modifying proteins, a Dam-X fusion protein modifies interacting DNA segments by methylating GATC stretches (G^me^ATC). This modification can be used to isolate interacting DNA regions by cutting with the methyl-specific restriction enzyme, *Dpn*I. In a typical DamID experiment, these fragments are amplified and subjected to DNA sequencing to generate genomic maps of interacting regions. The m6A-tracer system is an adaptation of the DamID technology for visualizing LADs in live-cell imaging [16]. Our variation of the m6A-tracer system relies on the demarcation of LADs by an inducible Dam-LMNB1 (DD-Dam-LMNB, Figure 1B) which methylates the lamina associated chromatin domains (LADs) underlying the nuclear lamina. Detection of the G^me^ATC-marked DNA (the LADs) is performed by the m6A-tracer, a catalytically inactive G^me^ATC binding domain of the *Dpnī* enzyme, fused to a fluorescent protein (such as mCherry).

To map the local proteome of the INM and LADs we generated 3 independent cell lines expressing either BirA*-Lap2β (to map the INM/lamina), BirA*-m6A-tracer alone (control), or BirA*-m6A-tracer with DD-Dam-LMNB1 (to map the LAD proteome) (Figure 1D). These cells were grown in the presence of exogenous biotin to enable efficient labeling of proximal proteins by BirA*. Cells harboring both BirA*-m6A-tracer and DD-Dam-LMNB1 constructs were split into two sets of cultured cells, one in the presence of the shield ligand and one without, as an additional control for the LAD proteome experiments (Figure 1D). In order to remove cytoplasmic contamination and limit our interrogation to the nucleus, a crude nuclear extraction preparation was performed on all cells. To detect biotinylation on candidate proteins we employ Biotinylation Site Identification Technology (BioSITe) and liquid chromatography-tandem mass spectrometry (LC-MS/MS) (Figure 1D) [31].

### Analysis of the LAP2β Interactome using BioID

In order to analyze the interactome of LAP2β, we used BirA* tagged LAP2β (BirA*-Lap2β) containing cells (Figure 1D). As a background control we used the BirA*-m6A-tracer construct expressed alone, without the presence of the DD-Dam-LMNB1 construct, which resulted in diffuse nuclear localization (red signal, Figure 1B) and subsequently similar biotinylation pattern (green signal, Figure 1B). A previous study utilizing LAP2β tagged with BirA* found that the expressed protein was mis-localized to the ER and cytoplasm. This is not very surprising since it has been previously shown that over-expression of LEM domain proteins leads to their accumulation in the ER and cytoplasm, and such mis-localization occurs with disease variants of lamin proteins as well [29,32,33]. We therefore sought to express our protein at low levels in order to preserve normal localization (Supplementary Figure 1). In addition, this low expression ensures minimal disruption to normal function of Lap2β and its partners and minimizes cellular stress due to overexpression (and mislocalization). In order to detect where our low expressing BirA*-Lap2β construct localized, we measured biotinylation signals using immunofluorescence on fixed MEFs expressing BirA*-Lap2β. As evidenced by the strong nuclear rim staining, we did not observe any gross mislocalization of LAP2β (Figure 1B). Some evidence of ER and mitochondrial expression is evident by the streptavidin signals, and this is to be expected since LAP2β transits the ER and there are endogenously biotinylated proteins in mitochondria. Therefore, as an additional step to maximize bonafide nuclear interactions, we performed a nuclear extract prior to protein isolation. While this extra step may result in the loss of some true nucleoskeletal-cytoskeletal interactions, it increases the rigor in detecting nuclear interactions. The correct localization of our LAP2β-BirA* (Figure 1B), its low expression (1-2% of endogenous levels of LAP2β, Figure S1) that obviates perturbations to the INM due to overexpression of a key INM protein, and a nuclear isolation step allowed for enriched detection of Lap2β-specific nuclear interactions.

Using BioSITe to detect biotinylated proximal proteins in our BioID screen we identified 334 total biotinylation sites in the BirA*-LAP2β containing cells and 684 sites in the BirA*-m6A-tracer alone containing cells (control). MS1 level quantitation was applied to obtain relative abundance differences between duplicate LC-MS/MS analyses of BirA*-LAP2β and BirA*-m6A-tracer cells. Replicate agreement was plotted (Figure 2A) and proteins enriched >2-fold were considered for LAP2β proximity/interaction (Table S1). Among the proteins enriched over control were expected and known interactors of LAP2β: LEM-domain containing protein 3 (MAN1), Emerin (EMD), and Lamins A and B1/2. Other known nuclear lamina proteins that were biotinylated were SUN domain contain proteins 1 and 2 (SUN1 and SUN2), lamin B receptor (LBR), and many nuclear pore complex members [4,34].

**Figure 2:**
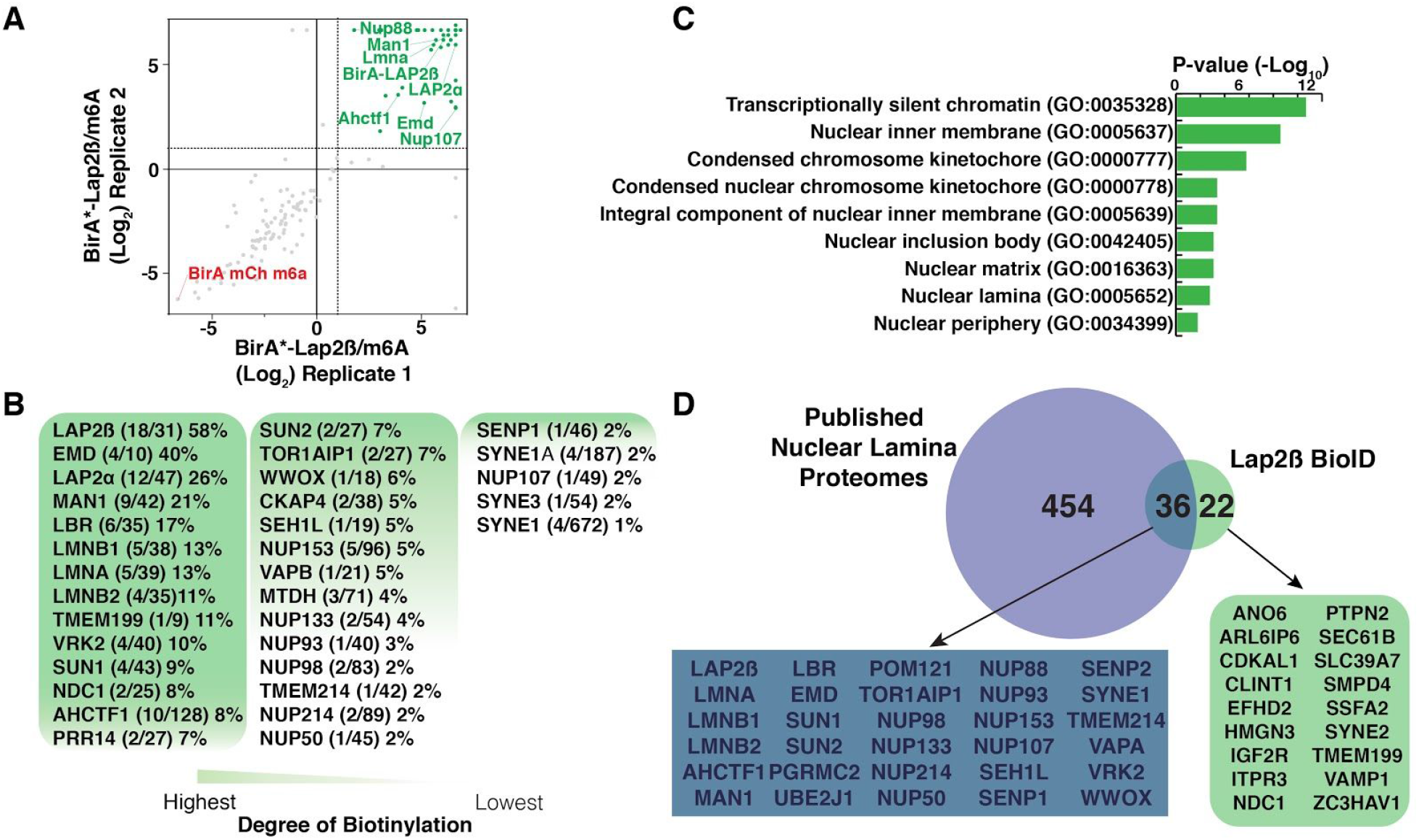
LAP2β BioID interactome. (A) Plot of replicate runs of MS1 level quantitation ratios of mass spectrometry identities of BirA*-LAP2β over BirA*-m6A-tracer alone containing cells. (B) Degree of biotinylation analysis of LAP2β interactome study. (C) Gene set enrichment analysis of LAP2β interactome. (D) Venn diagram of current LAP2β BioID interactome analysis and published nuclear lamina proteome analyses.

In an effort to rank the biotinylated proteins enriched over the control, in terms of potential closer proximity, we calculated the degree of biotinylation (number of biotinylated lysines/total lysines) and ordered proteins from greatest to least (Figure 2B) [31]. We detected that the bait LAP2β has the greatest degree of biotinylation followed by Emerin, MAN1, LBR, LMNB1/2, LMNA, and SUN1/2. We hypothesize that these proteins represent the most proximal and/or abundant interactors of LAP2β on the nucleoplasmic side of the INM.

To verify that our list of proteins was in fact enriching for nuclear envelope proteins we also submitted detected biotinylated proteins to cell component analysis using the Enrichr web based analysis tool [35]. Some of the top cellular component terms included nuclear periphery (GO:0034399), nuclear lamina (GO:0005652), and nuclear matrix (GO:0016363) (Figure 2C). To further examine if our data was consistent with previous studies we benchmarked our data on four published studies of nuclear envelope interactomes [21–23,27,28,36,37]: (*1*) Bar et al. combined a novel antibody based proximity labeling strategy and a meta-analysis of many lamin A and B interactome studies that included methods such as BioID, yeast two hybrid, and co-immunoprecipitation. (*2*) Kim et al. used BioID on many nuclear pore complex (NPC) members to build out the proximal proteins within the NPC and (*3*) Xie et al. used BioID on both A-type lamins. (*4*) Birendra et al. also used BioID to examine the interactomes of Lamin A and Sun2. We found 36 proteins to be overlapping between this list and our LAP2β interactome analysis (Figure 2D), including most of our proteins showing the highest degrees of biotinylation. We believe that these proteins represent very high confidence nuclear envelope proteins.

We do however note the slight representation of ER proteins and NUPs in our LAP2β dataset which includes vesicle-associated membrane protein-associated protein B (VAPB) and nucleoporin NDC1 (NDC1). We hypothesize that this could be due to the NPC presenting itself as a kinetic bottle-neck for the relatively large BirA*LAP2β and therefore, a longer residence times in the ER and NPCs. Additionally, notably absent from our Lap2β interactome analysis is barrier to autointegration factor (BAF) which is a known interactor of Lap2β, as well as HDAC3, another putative interactor [38–40]. We hypothesize that the small size of BAF (~10 kD) makes detection of this protein by mass spectrometry based approaches particularly challenging. It is also important to emphasize that BioID is a label transfer technique and hence, the biotinylation of potentially transient interactors such as HDAC3, coupled with limited material due to the sub-physiological expression level of BirA*-LAP2β, it is possible that some of these might have been missed by our analysis.

Given that LAP2β contains a transmembrane domain which is localized within the INM, we next examined our data for transmembrane domain containing proteins. We observed that 34 of the 60 LAP2β proximal proteins contained a transmembrane domain (Table S1). It has been shown that membrane topology can be predicted with biotinylation site level data [31]. Many of these proteins exhibited biotinylation on areas of the protein consistent with lumen/nucleoplasmic annotated topology (Table S1). Interestingly, 18 of these proteins also have subcellular annotation for the endoplasmic reticulum (ER) including VAPB, CKAP, and TMEM214 which could indicate unknown nuclear localization and function, however it is also possible that these proteins are part of the ER trafficking of LAP2β to the nucleus [41].

### LAD directed BioID using m6A tracer system

Having identified and refined the proteome at the lamina, we next asked if we could discriminate proteins that are proximal to LADs and their relationship with the nuclear lamina network using our m6A tracer system which is built off the sequencing technique DamID. In DamID, DNA Adenine Methyltransferase (Dam) derived from *E. coli* is fused to a DNA interacting protein [9]. Dam adds methyl groups to adenines in the vicinity of the fusion protein, thereby marking the interaction sites on the proximal DNA segments (G^me^ATC) which can be cut with the G^me^ATC specific restriction enzyme *Dpn*I. Fragments are then subjected to ligation mediated PCR and deep sequencing. Using Dam-LaminB1 and Dam-only (normalizing control) to mark DNA, the resulting data identify regions of DNA associating with the nuclear lamina, Lamina Associated Domains (LADs), expressed as a relative enrichment ratio: log_2_ (Dam-Lamin/Dam only). The m6A-tracer system is an adaptation of the DamID technology for visualizing LADs in single-cell live imaging [16]. This system relies on the demarcation of LADs by an inducible Dam-LMNB1 (in this study, DD-Dam-LMNB1), the expression of which can be exogenously controlled via Shield1 ligand masking of the destabilization domain, and detection of this marked DNA (the LADs) is performed by the m6A-tracer, a catalytically inactive G^me^ATC binding domain of the *Dpn*I enzyme, fused to a fluorescent protein (BirA*-m6A-tracer in Figure 1B) [42–45]. The DD domain prevents accumulation of the fusion protein via proteasomal degradation in the absence of shield ligand (Figure 1B-D) [46,47]. Upon introduction of shield ligand, the protein is stabilized, enabling the DD-Dam-LMNB1 protein to methylate LADs.

In order to identify LAD-proximal proteins, we compared biotinylated peptides from cells harboring DD-Dam-LMNB1 and BirA*-m6A-tracer with (marks LAD-proximal proteins) or without (background control) shield ligand. LAD-specific interactions were determined to be >1.6-fold average ratio between replicates (replicate agreement is plotted in Figure 3A) of plus over minus shield ligand (i.e recruited to LADs over background). Using BioSITe to detect proximal proteins, we found 1179 biotinylation sites in the plus shield condition and 1128 in the minus (Table S2) [31]. Using this approach to map the proteome of LADs, we were able to identify three major classes of proteins enriched in the plus shield condition: INM proteins, cell cycle related and DNA binding/chromatin proteins (Figure 3B, C).

**Figure 3:**
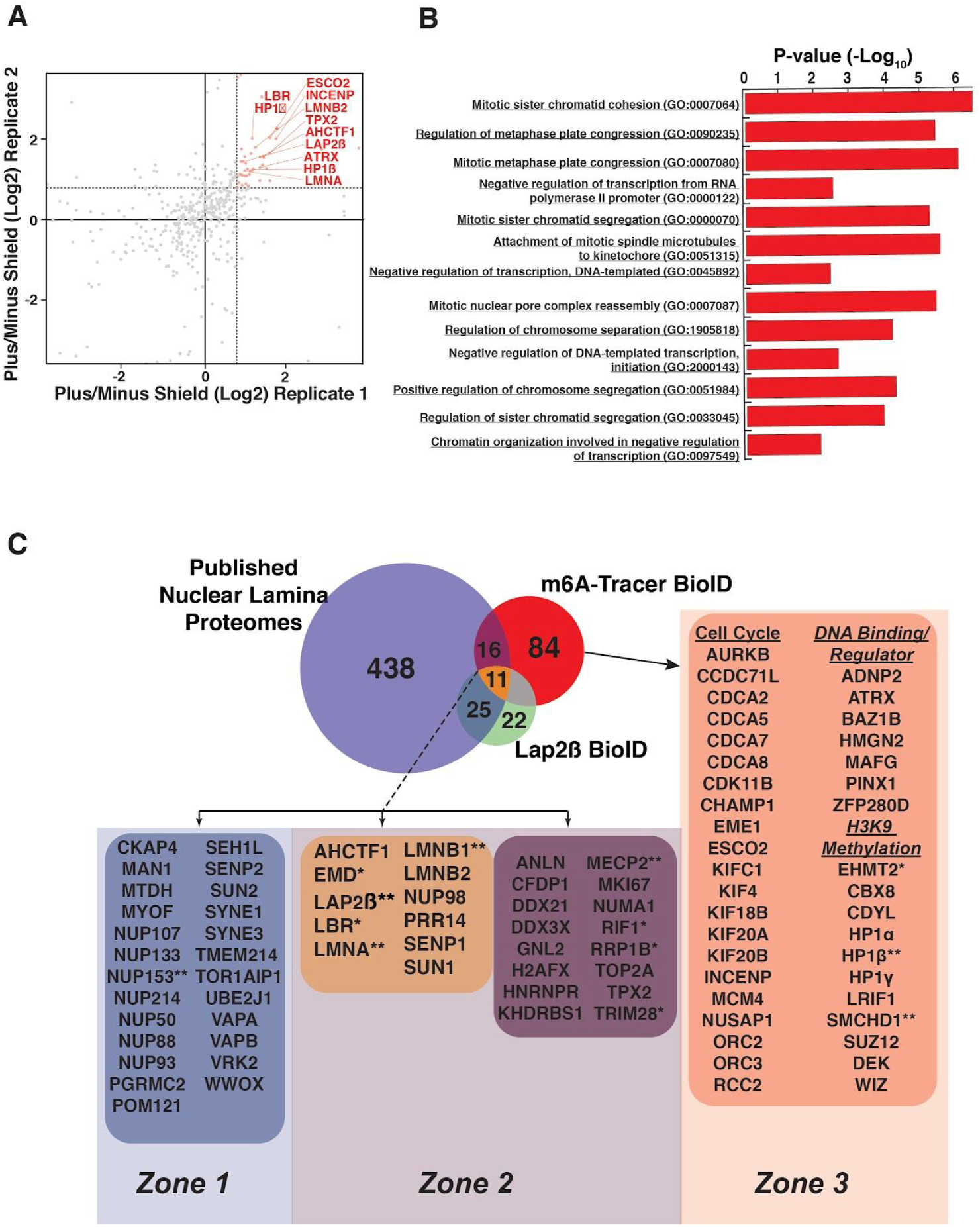
LAD microproteome analysis. (A) Plot of replicate runs of MS1 level quantitation ratios of mass spectrometry identities of BirA*-m6A-tracer/DD-Dam-LMNB1 plus shield over minus shield ligand. (B) Gene set enrichment analysis of m6A-tracer BioID LAD proteome. (C) Integrative venn diagram of published nuclear lamina proteomic analyses, our current LAP2β BioID interactome analysis and the LAD-ome analysis. Proteins are further classified into zones depending on whether they solely interact with INM/lamina components and not LADs, (zone 1), interacted with both LADs and other INM/lamina components (zone 2) or are LAD specific (zone 3). Proteins marked * have been validated by various groups and the experiments and results are summarised in Table 1. Proteins marked ** have been bioinformatically validated in house.

Specifically, to better understand the 111 LAD-proximal proteins that were enriched over our minus shield control, we submitted these proteins to pathway analysis using the Enrichr web-based pathway analysis (Figure 3B) [35]. Of the most significantly enriched biological processes, many were processes in negative regulation of DNA transcription and cell cycle related pathways. Examples of the latter include: mitotic sister chromatid cohesion and segregation, regulation of mitotic metaphase plate congression. Notable cell cycle related genes include many members of the cell division cycle associated protein family including 2 (CDCA2), 5 (CDCA5), 7 (CDCA7), and 8 (CDCA8), many kinesin family members such as 4A (KIF4A), 18B (KIF18B), 20A (KIF20A) and 20B (KIF20B). We also observed that many microtubule/spindle attachment proteins related to the kinetochore including the serine/threonine kinase that regulates segregation of chromosomes during mitosis Aurora kinase B (AurkB), nucleolar and spindle associated protein 1 (Nusap1), regulator of chromosome condensation 2 (RCC2), microtubule nucleation factor (TPX2), sister chromatid cohesion protein (PDS5) and inner centromere protein (INCENP). In addition, the replication timing factor Rif1, was identified as LAD-interacting, in agreement with previous studies [48,49]

As expected, another highly represented class of proteins enriched on LADs were proteins of the INM. These included many known INM proteins such as LAP2β, MAN1, Emerin, Lamins A and B1/2, LBR, and SUN1 (Figure 3C). Strikingly, NUP proteins were not enriched in the LAD proteome, suggesting that these chromatin domains are indeed not proximal to nuclear pore complexes, in agreement with cytological data.

### LADs are particularly enriched in chromatin modifying and binding proteins

As mentioned above, one of the major classes of proteins enriched on LADs were involved in negative regulation of transcription. Specifically, many of the identified proteins have roles in the establishment/regulation of H3K9 methylation or bind to H3K9 methylated histones, such as heterochromatin protein 1 alpha and beta (HP1α. HP1β), polycomb repressive complex 2 (PRC2) subunit Suz12, TRIM28 (Kap1), SMCHD1, PRR14 and DEK (Figure 3C). Of particular interest were the abundance of LAD-interacting proteins involved in establishment and maintenance of heterochromatin, specifically euchromatic lysine methyltransferase 2 (EMHT2) and its binding partners chromodomain Y like (CDYL) and the transcription factor widely interspaced zinc finger motifs (WIZ). These proteins have been shown to facilitate mono- and dimethylation of H3K9.

Finally, we also observed many additional chromatin modifying and DNA binding proteins such as AT-hook containing transcription factor 1 (AHCTF1, also known as ELYS, which has also been implicated as a bona fide NPC protein [50,51]), ATRX chromatin remodeler (ATRX), MECP2, tyrosine-protein kinase BAZ1B (BAZ1B), high mobility group nucleosomal binding domain 2 (HMGN2), and PIN2/TERF1 interacting telomerase inhibitor 1 (PINX1) [50,52–55]. We also identified a few transcription factors not known to be involved in heterochromatin such as MAF BZIP transcription factor G (MAFG), zinc finger protein 280D (Zfp280d), and ADNP homeobox 2 (ADNP2).

### Integration of laminome and LADome data identify unique and overlapping micro-proteomes

We next sought to determine the overlapping proteomes between the nuclear lamina and LADs. We focused on using existing data and our new Lap2*β* proteome to maximize potential proteins interacting with chromatin. There were 52 proteins that overlapped between the published laminome data and our Lap2*β* proteome or the m6A-tracer LAD proteome (Figure 3C). Importantly, the well known nuclear lamina proteins such as LBR, LMNA/C, LMNB1, and SUN1 all showed enrichment on LADs as well (Figure 3C, zone 2). In contrast, while the published laminome and Lap2*β* showed interactions with nuclear pore complexes (NUPs, Figure 3C, zone 1), these were largely missing from the LAD interactome data. This finding supports cytological and DamID studies suggesting that the chromatin underlying the NUPs is more euchromatic and distinct from the peripheral heterochromatin that comprises LADs.

In total, we identified 27 proteins, including MECP2, AHCTF1 (also known as ELYS), and PRR14, that were found to be biotinylated in the BirA*-m6A-tracer experiment that were also detected in the published nuclear lamina interactomes and/or our Lap2*β* interactome (Figure 3C, zone 2). These are potential candidates for proteins that are at the interface of the INM and LADs, perhaps linking them together and regulating dynamic LAD organization [4].

The 47 proteins detected only in the LAP2β-BioID experiment, as well as the majority of proteins in the published laminome data that are not LAD-enriched (Figure 3C, zone 1), likely represent more inner nuclear membrane proximal proteins that do not interact with and are farther from the underlying chromatin (LADs). These include the previously described NUP proteins. Other examples include, Progesterone Receptor Membrane Component 2 (PGRMC2), Torisin-1A-interacting protein 1 (Tor1aip1), Myoferlin (MYOF), transmembrane protein 214 (TMEM214), metadherin (MTDH also called protein LYRIC), cytoskeleton associated protein 4 (CKAP4), vesicle-associated membrane protein-associated protein B/C (VAPB); these are all transmembrane containing proteins that may not have large enough nucleoplasmic domains to be in proximity to the LADs, and, for the Lap2*β*-specific interacting transmembrane proteins, could also represent transient interactions from transit through the ER. Nonetheless, these proteins appear to be more distal to the underlying LAD chromatin.

84 proteins were identified as associating *uniquely* with LADs (Figure 3C, zone 3). Since the LADs and the lamina are in such close proximity, we were surprised to find such a clear distinction between the LADome and the laminome. The LAD-specific proteins include the previously described chromatin modifier complexes such as EHMT1/2, cell cycle regulators and chromatin interactors. This is particularly interesting given the recent evidence that chromatin state directs LAD organization [12,14–18]. These data suggest, and support previous findings, that these regions are enriched for chromatin complexes that initiate and maintain a heterochromatic state [10,12]. These data also support a model in which chromatin state is established independently from association of these chromatin domains with the lamina, since these proteins do not interact with the INM/lamin directly [15,17].

### Validation and other supporting data

To our knowledge, this presents the first comprehensive characterization of the local proteome associated with LADs. Many of the proteins identified have already been extensively validated, with some of these validations corroborating the placement of the proteins in the specific peripheral zones. Published and new validation experiments and data that directly show association with the INM/lamina and/or involvement in LAD establishment/maintenance have been summarised and compiled in Table 1 [10,12,16–19,48,49,56–67].

**Table 1.**
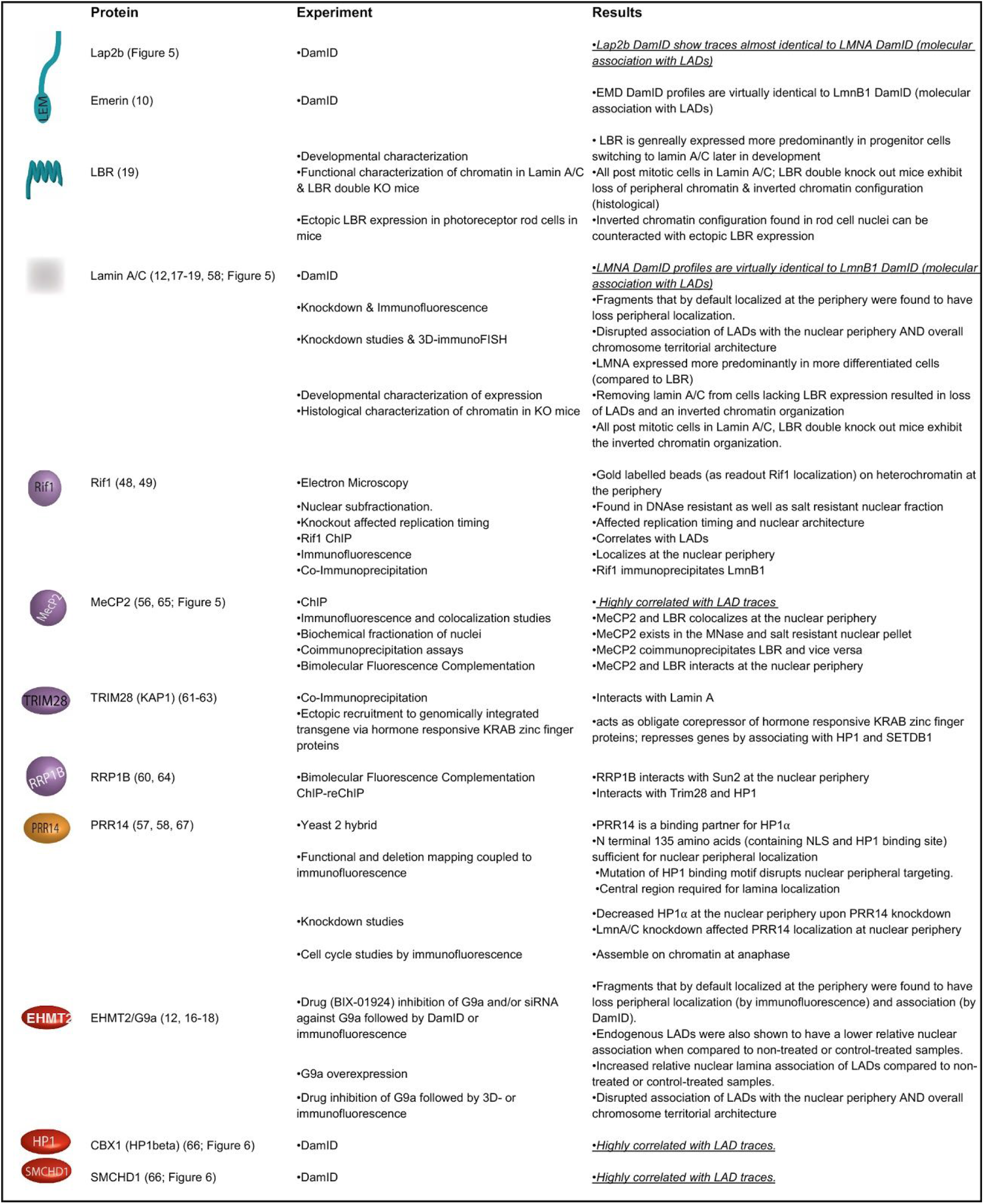
(10, 12, 16–19, 48, 49, 56–67): Table detailing experiments and results from published work directly validating protein hits from our BioID study.

#### Zone 1 Proteins at the INM/lamina that do not interact with LADs

As previously mentioned, proteins classified in zone 1 likely represent nuclear envelope proteins that are farther from and do not interact with LADs. To test this supposition, we interrogated the interaction of one such protein, Nup153, with LADs using publicly available data (Figure 4, [10,68]. Nup153 has previously been shown to interact with chromatin, but our data suggests that this protein is excluded from LAD chromatin domains. We find that the majority of Nup153 interactions with chromatin occur outside of LADs and, intriguingly, Nup153 peaks that appear to be within LADs (at a gross scale) coincide with regions that have low lamin B1 signal, which we have previously termed DiPs (Depleted in Peripheral signal, [18]; Figure 4). Interestingly, these DiPs correlate with active transcription start sites and enhancers, with a distinct and discrete switch from the inactive B to the active A chromatin compartment, These combined data support our integrated BioID data which suggests that the Nup153 interactions with chromatin at the lamina are more distal to LADs (Figure 4).

**Figure 4:**
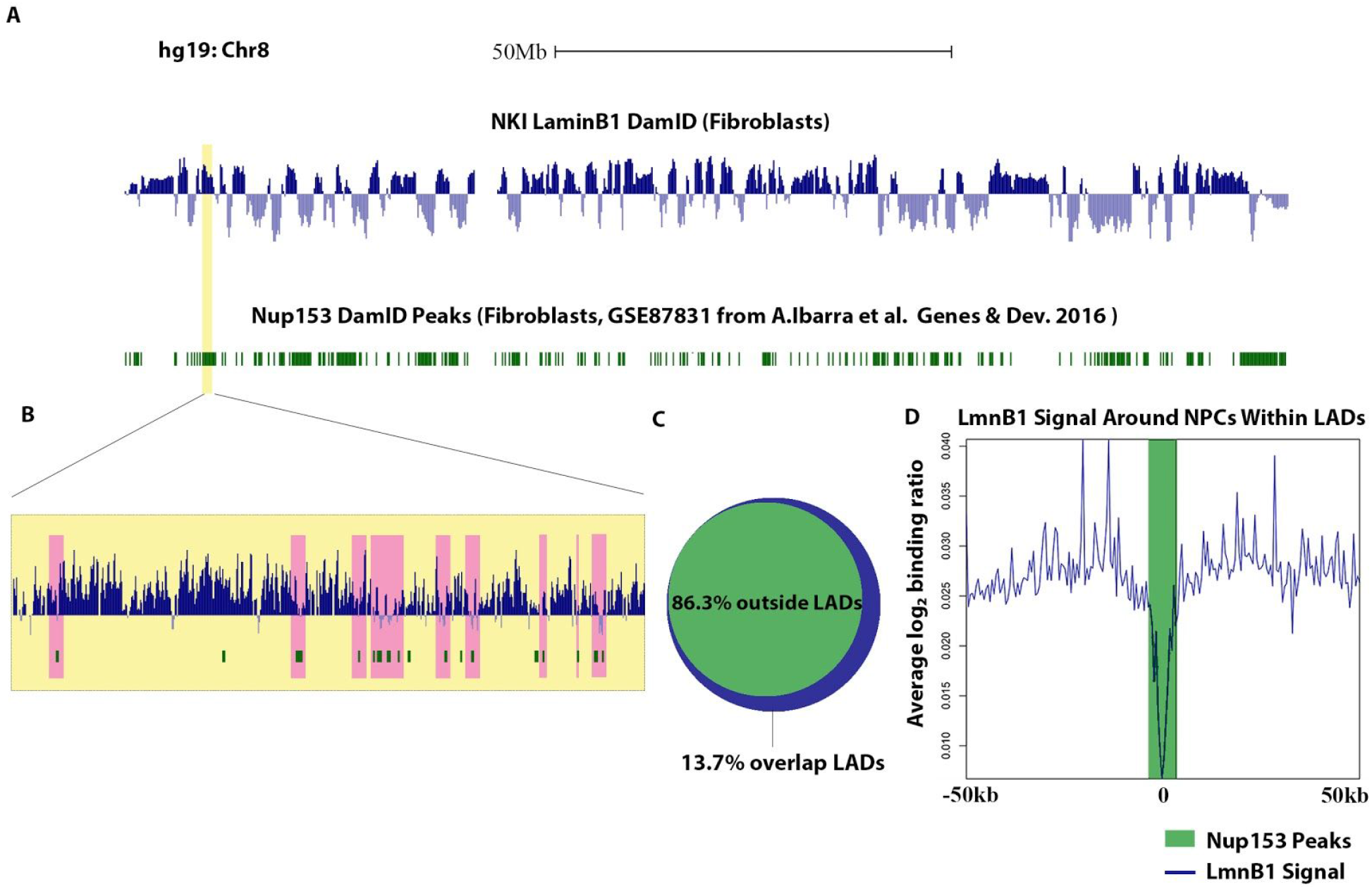
(A) Log2 ratio plots from human fibroblasts (hg19 build) chr 8 of LmnB1 DamID (blue) and Nup153 DamID peaks in green[10,68]. (B) Inset shows a magnified view of a LAD that appears to be highly dense with Nup153 binding sites. (C) Venn diagram showing the percentage (in base coverage) of Nup153 distribution relative to LADs. (D) LmnB1 profile anchored at all LAD-Nup153 peaks (the 13.7% that are found within LADs) centers. Line graphs represent the average trend across all such Nup153 peaks.

#### Zone 2 Proteins that bind the INM/Lamina network and LADs-the “middlemen”

Proteins classified in this zone represent candidates that interface the INM and LADs, perhaps linking them together and regulating dynamic LAD organization [4]. This zone is comprised of both INM/Lamina proteins and chromatin binding proteins. It is hardly a surprise that lamin proteins fall into this category as they have been shown to interact with LADs and to be important for their localization to the nuclear periphery (Table 1 [10,12,16–19,48,49,56–67]; Figure 5). In addition to the lamin proteins, some INM proteins were also identified as spanning the INM/lamina and interacting with chromatin. LBR has previously been shown to interact with chromatin and is required for normal chromatin domain organization (Table 1 [10,12,16–19,48,49,56–67]) Similarly, Emerin interactions with chromatin, as measured by DamID, are highly correlated with LADs [10]. In addition, both LBR and emerin have been identified in multiple BioID and biochemical experiments to interact with the same lamin network [10]. These combined data support our findings here that these proteins span the lamina network and chromatin interface. Similarly, we find that Lap2*β* itself also falls into this category. Lap2*β* has been shown, through multiple experiments, to interact with the lamina and INM network. In addition, several studies have suggested that this protein is important for LAD organization and regulation[12,69]. We decided to interrogate the interaction of Lap2*β* with LADs to verify that this protein indeed interacts across the INM/chromatin interface, as our BioID data suggests. Using a DamID approach, we find that Lap2*β* interaction with chromatin occurs on LADs; the Lap2b interaction signatures are virtually identical to LADs. (Figure 5 A-B and Figure S2).

**Figure 5:**
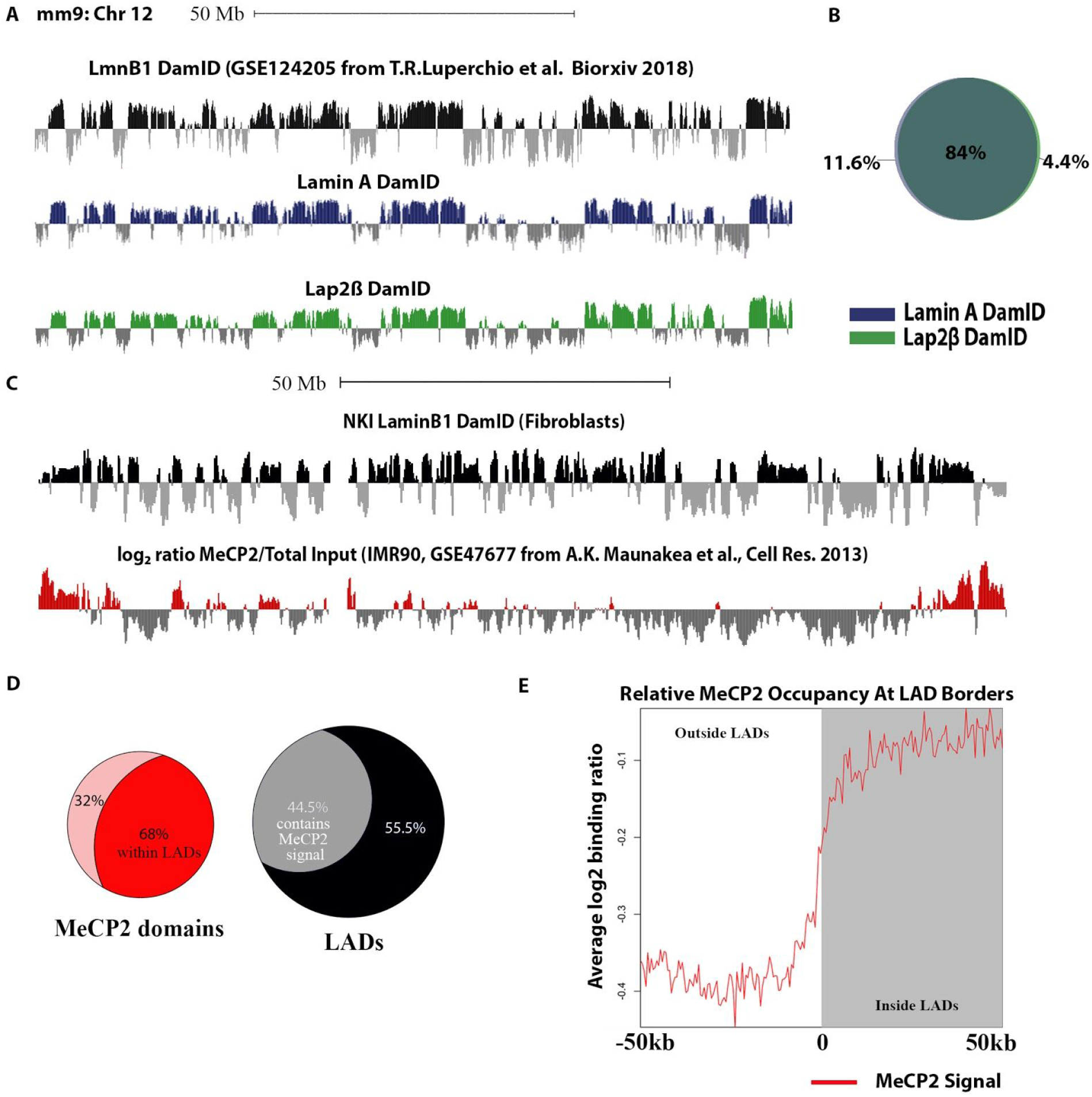
(A) Log2 ratio plots from MEFs (mm9 build) chr 12 of LmnB1 DamID (black), our LaminA (blue) and Lap2β DamID (green)[17]. (B) Venn diagram showing degree of overlap in percentage between lamin A and Lap2β LADs. (C) Log2 ratio plots from human fibroblasts (hg19 build) chr 8 of MeCP2 occupancy (ChIP) in red, LmnB1 DamID in black[10,65].(D) Venn diagrams showing the percentage (in terms of base coverage) of MeCP2 domains within LADs and the percentage (in base coverage) of LADs that are bound by MeCP2 (E) MeCP2 profile anchored at all boundaries of LADs of size 100 kb or greater and oriented from outside LAD (left) to inside LAD (right). Line graphs represent the average trend across all boundary profiles.

In addition to well known resident proteins of the INM/lamina, other proteins less often associated with INM/lamina studies were also classified in this zone as potential INM/LAD links. An example would be methyl-CpG binding protein (MECP2), which we have found to interact with both LADs and in published laminome data. MECP2 is a known repressor of DNA that binds to methylated CpG regions in the DNA. Recently, a study demonstrated that CpG methylation demarcates the heterochromatic “B-compartment” of the genome, which is highly correlated with LADs [17,18,70]. Consistently, bioinformatic analysis of publicly available MeCP2 ChIP-seq data shows a high coincidence of MeCP2 binding domains with LADs - with two thirds of MeCP2 occupancy coinciding with LADs and almost half of the LADs potentially regulated by MeCP2 binding events (Figure 5 C-E [10,65]). Furthermore, an interaction between LBR and MECP2, as well as HP1, has been observed, further implicating MECP2 physically interacting with the INM proteome and the LAD proteome [56,71–75]. Other examples in this category include: TRIM28, RRP1B, which have both been documented to interact with nuclear lamina components and shown to be associated with H3K9 methylated locations in the genome; PRR14, which has been shown to provide a link between LADs and the INM through its association with HP1; and RIF1, a known chromatin modifier involved in replication timing and previously shown to be enriched on LADs (Table 1 [10,12,16–19,48,49,56–67])

#### Zone 3 proteins associating uniquely with LADs

Lastly, proteins classified in this zone potentially regulate/modulate LADs independent of their association with the nuclear periphery. Proteins in this zone fall under 3 sub-categories comprising of cell cycle regulators, proteins that bind and/or regulate DNA and proteins bind to or modulate H3K9 methylation. We were particularly interested in the proteins that affect H3K9 methylation since the methylation of H3K9 is an important feature of heterochromatin and has been shown to be enriched at and required for LADs [10,11,15,76–79]. This includes the HP1 proteins, chromobox 5 (CBX5) also known as heterochromatin protein 1 alpha (HP1α) which binds to H3K9me3 [80] and heterochromatin protein 1 beta (HP1β or CBX1). CBX1 serves a similar function as HP1α and, as shown in our bioinformatic analysis, to be highly coincident with LADs (Table 1 [10,12,16–19,48,49,56–67], Figure 6 A, B, C [66]). We find the 80% of CBX1 binding peaks reside within LADs (Figure 6 B). Additionally, we detected other chromobox family proteins: chromobox 3 (CBX3 or HP1γ) and 8 (CBX8) in our BioSITe data. HP1 proteins, particularly HP1α, bind to methylated H3K9 and mediate gene silencing by maintaining and spreading heterochromatin. Alongside these HP1 proteins we also identified the histone chaperone and proto-oncogene DEK, which, has been shown to prevent access to transcription machinery and to interact with HP1, enhancing its binding to H3K9me3 [82], [83]). Furthermore we found the SUZ12 polycomb repressive complex 2 subunit (Suz12) which has been found to both stabilize HP1α and increase H3K27me3, which in turn increases the HP1α/β/γ binding of H3K9me3[84].

**Figure 6:**
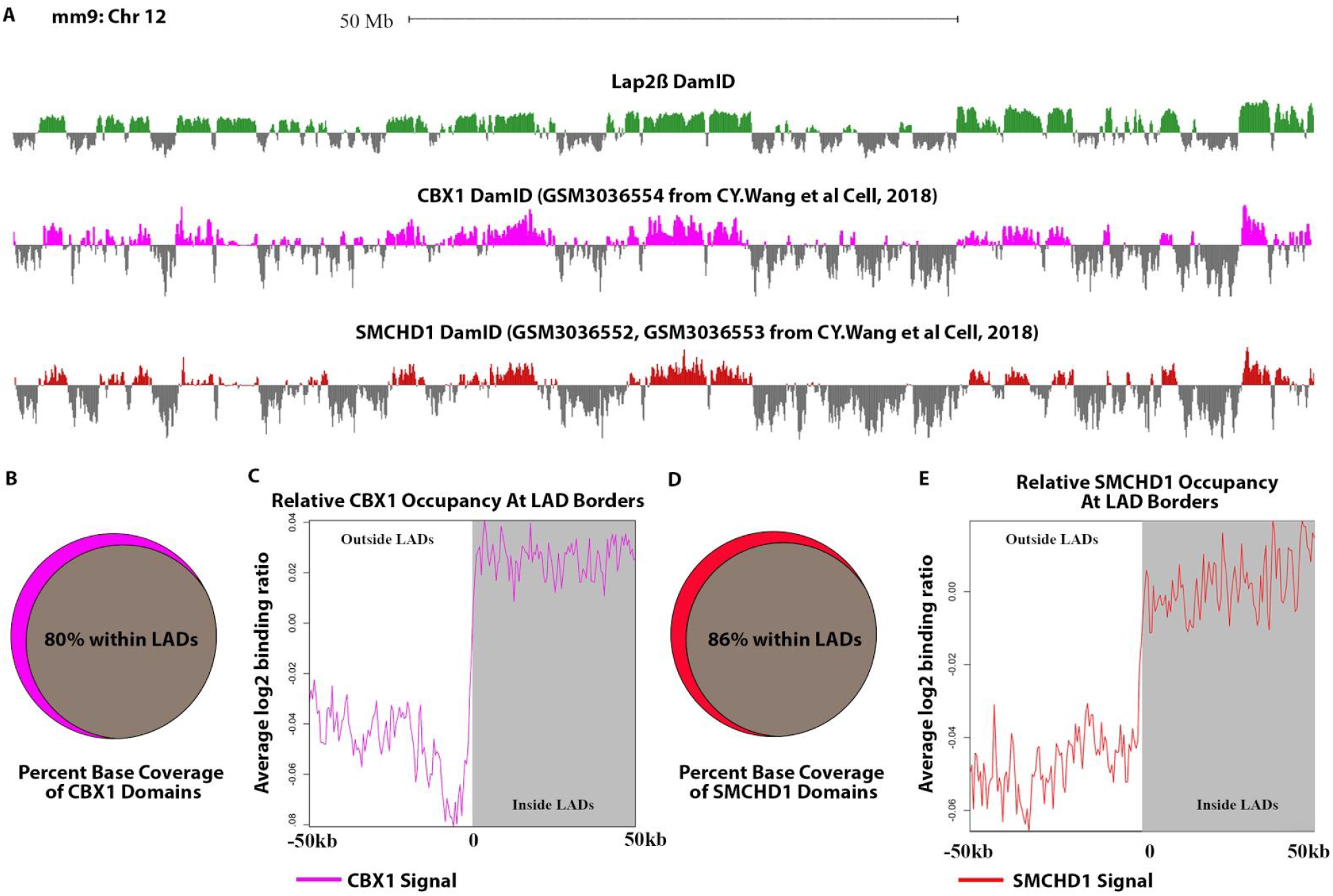
(A) Log2 ratio plots from MEFs of our Lap2β DamID (green), CBX1 (pink) and SMCHD1 DamID (Red)[66]. (B) Venn diagram showing percentage (in base coverage) of CBX1 occupied genomic domains that overlap LADs. (C) CBX1 profile anchored at all boundaries of LADs of size 100 kb or greater and oriented from outside LAD (left) to inside LAD (right). Line graphs represent the average trend across all boundary profiles for each feature. (D)Venn diagram showing the percentage (in base coverage) of SMCHD1 occupied genomic domains that overlap LADs. (E)SMCHD1 profile anchored at all boundaries of LADs of size 100 kb or greater and oriented from outside LAD (left) to inside LAD (right). Line graphs represent the average trend across all boundary profiles for each feature.

Another intriguing protein that we identified in this zone is the euchromatic lysine methyltransferase 2 (EHMT2, or often called G9a). EHMT2 is an important methyltransferase known to methylate H3K9 and H3K27, facilitating H3K9me1 and H3K9me2 in particular, and subsequent gene silencing [85]. Disruption of this protein by either epigenetic drugs or shRNA-mediated silencing disrupts peripheral heterochromatin and LAD formation (Table 1 [10,12,16–19,48,49,56–67]. Intriguingly, this protein was also recently implicated in methylating LaminB1, an event suggested to be important for LAD organization (although, we note, that we did not detect interactions of LaminB1 and EHMT1/2 across multiple datasets) [87]. Moreover, EHMT2 was found to interact with and is recruited by chromodomain Y like (CDYL), another protein we identified to be enriched at LADs, to facilitate H3K9 dimethylation and repression of the neurogenesis master regulator, REST, target genes [88,89]. EHMT2 and EHMT1 have been found to form a complex with widely interspaced zinc finger motifs (WIZ), another protein we found in our data, which acts to stabilize this complex on chromatin and prevent degradation of EHMT1 [90,91].

Interestingly, we also identified structural maintenance of chromosomes flexible hinge domain containing 1 (SMCHD1) protein as LAD enriched--a protein that had not been previously identified on LADs. On inactive X chromosomes, SMCHD1 and HP1 are found to colocalize at areas of H3K9 methylation [66,81,92]. SMCHD1 has also been shown, by co-immunoprecipitation, to interact with ligand dependent nuclear receptor interacting factor 1 (LRIF1), another protein identified in this study, which helps target the protein to H3K9me3 regions [93]. In order to test whether SMCHD1 interacts with LADs, as indicated by our proteomic data, we compared published SMCHD1 DamID data (from a study focused on the structural regulation of the inactive X chromosome and not LADs) with our Lap2β DamID data. This analysis showed a remarkably high coincidence rate of 86%, of SMCHD1 binding domains with LADs suggesting that SMCHD1 potentially regulates LADs on top of its known regulatory roles in establishing and/or maintaining the inactive X chromosome (Figure 6 A, D, E [66]).

## Conclusions and discussion

It remains unclear how lamin associated domains (LADs) are organized and maintained at the nuclear lamina. Our previous studies had implicated Lap2β in LAD organization, but DamID maps of DNA interactions with this protein were indistinguishable from LADs identified using LaminA or LaminB, both of which have been used previously to demarcate these DNA domains (Figure 5, Figure S2) [18]. In addition, previous studies have shown that both chromatin state of the LADs and Lamin A/C are critical for LAD organization, suggesting the potential for chromatin scaffolding protein “middlemen” in mediating interactions between LADs and the lamina [94]. In support of this idea, PRR14, a protein that interacts with HP1α (heterochromatin protein 1α), has been shown to interact with both the lamina and chromatin [95]. In order to elucidate potential mediators of LAD organization, we therefore sought to map the overlapping proteomes of LADs and the nuclear lamina (Figure 1).

Previous studies have identified proteins enriched at the nuclear lamina (the “laminome”) through multiple methods, including BioID [21–25]. Here we use BioID with BioSITe (BioID coupled with biotinylation site enrichment and analysis) to measure protein-protein interactions of the INM protein Lap2β (Figure 2) in MEFs and integrate these with the previous laminome findings to generate an enhanced INM/lamina proteome map [21–23,27,28,36,37]. Not surprisingly, the majority of Lap2β protein-protein interactions uncovered in this study were previously identified as part of the laminome. With few exceptions, the majority of proteins unique to Lap2β interactions are other transmembrane proteins, many of which have identified roles at the plasma or cytoplasmic membranes, likely reflecting transient interactions as Lap2β transits the ER/Golgi. Exceptions to this include HMGN1, a minor groove AT-hook DNA-binding protein, and Nesprin-1 (SYNE-1), a LINC-complex protein linking the cytoskeleton to the nuclear lamina network.

To find LAD-enriched proteins, we used our BioID with BioSITe pipeline coupled with LaminB1-directed DamID. We leveraged a modified version of a live cell LAD visualization system to recruit biotin ligase directly to LAD chromatin (Figure 1B-D). As with the Lap2β-directed BioID experiments, we verified that the biotin ligase marked proteins at the periphery of the nucleus (Fig 1B). LAD-enriched proteins included cell cycle related proteins, proteins of the INM/lamina and chromatin interactors and modifiers (Figure 3B, C). Strikingly, the majority of the proteins identified in the LADome were LAD-specific, that is to say, they were not identified in either previous LAD interactome data or in our Lap2β interacting protein set. Remarkably, LADs did not generally interact with NUPs, which is in agreement with cytological data and DamID data showing that NUPs interact primarily with transcriptionally active regions of the genome [96]. Given that NUP98, out of all the NUPs identified in the laminome, is the only NUP that was also identified as LAD-interacting possibly suggesting that NUP98 interacts with LAD border regions given that it has been shown to interact with highly transcribed genes and highly active genes that flank LADs, demarcating their borders [10].

The LAD-specific interactions highlight that, although LADs and the INM/Lamina are spatially proximal, they are indeed different, but overlapping, microenvironments (Figure 3 and 7). It is especially intriguing that a large fraction of the proteins that are identified as LAD-specific are related to modulating chromatin state, particularly H3K9me2/3. Recent data from our lab has shown that, through the cell cycle, LADs within a chromosome self-interact prior to their organization at the nuclear lamina, suggesting that the higher order interactions of these chromatin domains are independent of their lamina association [17]. However, the inactive chromatin state, particularly H3K9me2/3 and H3K27me3 are required for lamina association [12–15]. These data are compatible with a two-step model in which chromatin state mediates self-association of these domains and that lamina association is then mediated by proteins which interact with (and help maintain) these specific chromatin modifications or states.

**Figure 7:**
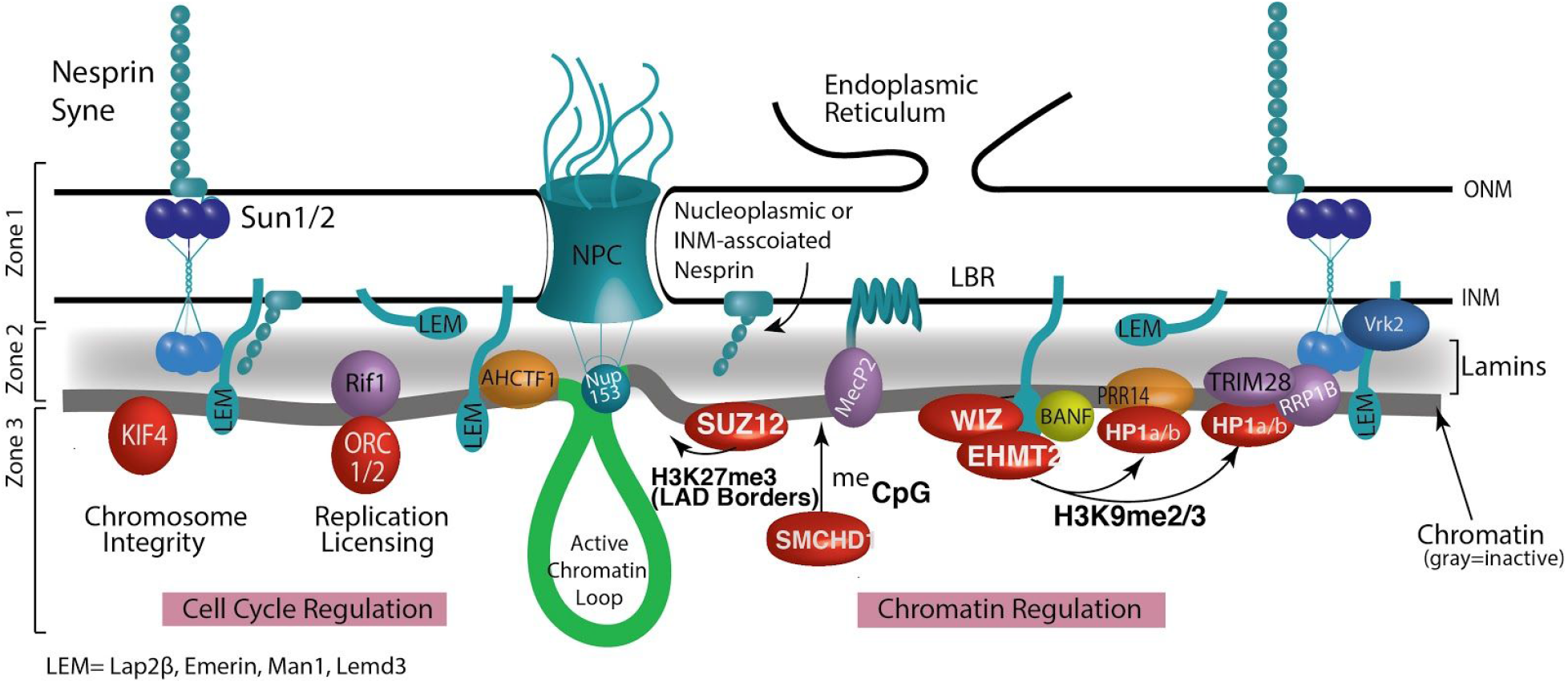
Model of the local proteome at the INM/lamina/LAD interface as resolved by BioID experiments. The model depicts the nuclear periphery resolved into 3 zones. Zone 1is the furthest away from chromatin comprising proteins that do not interact with LADs, such as NPCs, some nesprins, and regulators of LEM domain proteins. Zone 1 then transitions into Zone 2 comprising of lamin proteins (shaded gray) and membrane-spanning proteins (colored teal) that traverse the lamina/LAD interface. In this region, we also find a unique set of chromatin/DNA binding proteins that display interactions with both the INM/lamina network and LADs (purple and gold). BANF, while not found in our datasets, is included in this zone based upon numerous in vivo and in vitro studies demonstrating its position as a linker between chromatin and the nuclear envelope. Thus, Zone 2 is made up of a lamin scaffold, INM proteins, and a specific subset of DNA interacting proteins that facilitate LAD interactions with the nuclear lamina. Finally, Zone 3 shows LAD-specific proteins (red) identified in our screen. These are highly enriched in chromatin-binding and modifying proteins.

Of particular interest, then, are the proteins that sit at the interface of the INM/lamina and LADs. LADs were initially identified as domains of chromatin in molecular proximity to LaminB1, using the proximity labeling method DamID. These “middlemen” proteins, including Lamin A/C, emerin, and LBR, which have been previously shown to be necessary for LAD organization. In this and other studies, lamin A/C and emerin have also been used to identify LADs, thus their identification in our screen as overlapping LADs and the INM/lamina is not surprising [59,97,98] (Figure 3C and Figure 7, zone 2). These “middlemen” proteins are all potential mediators of LAD organization at the lamina [12,17–19]. Thus, our analysis has identified known mediators of LAD scaffolding as proximally interacting with LADs.

Importantly, our BioSITe analysis also uncovered additional proteins that sit at this interface and are, heretofore, untested potential mediators of LAD organization to the nuclear lamina (Figure 3C, zone 2).These include the previously implicated PRR14, as well as RIF1, AHCFT1, and MECP2 and H2AFX. RIF1 binding, which demarcates late replicating domains, has previously been identified as overlapping with LADs and in regulating genome organization [49]. AHCTF1 (also known as ELYS) is implicated in nucleopore complex assembly, proper exit from mitosis, and genome stability [50,51]. MECP2, a methyl CpG binding protein, has clear roles in establishing and maintaining heterochromatin. MeCP2 partners with HDACs and histone methyltransferase and is involved in higher order chromatin remodeling and silencing [52,99–105]. We have also shown bioinformatically using published MeCP2 ChIP datasets that MeCP2 is enriched within LADs (Figure 5). Intriguingly, while our proteomic analysis was not suited to identifying chromatin modifications, we identified histone H2AFX, which is phosphorylated to form γH2Ax. γH2AX is found on double strand breaks on chromatin, which become heterochromatinized and possibly shunted to the nuclear periphery [106]. H2Ax may also be key in establishing de novo LAD organization since one study found that during epithelial to mesenchymal transition, regions that demarcate newly formed LAD borders become enriched in γH2AX[107].

The synthesis of this data allows us to better refine a model of the micro-proteome of the INM/lamina and LAD interface (Figure 7). Taken together, our findings suggest that LADs and the INM/Lamina have distinct, but overlapping proteomes. Several of the proteins that are shared between LADs and the INM/lamina have already been shown to be important for organization of LADs to the lamina. We propose that several newly identified proteins spanning these two proteomic domains are likely involved in establishment or maintenance of LAD chromatin. Moreover, our findings are consistent with LADs supporting a heterochromatic architecture that is independent of their association with the nuclear lamina, but important for mediating interactions at the lamina.

## Acknowledgements

We thank Rakel Tryggvadottir and Sinan Ramazanoglu for invaluable help with sequencing. This work was funded in part by NIH grant R21AG050132. X. W. and V. H. were supported in part by NIH grant R21AG050132. V.H. and J.A. C. were supported in part by NIGMS Training Grant 5T32GM07814. M.G. was supported by NIH Training Grant T32GM007445.

## Methods

### Plasmid Construction

To clone Fu CMV DD-Dam-LMNB1, Dam LmnB1 was first amplified from Plgw Dam LmnB, kindly provided by the Van Steensel lab, with overhanging restriction sites. The amplicon was then TA cloned into pGEM-t easy (promega). The cdt1 fucci tag, flanked by *Age*I and *Xma*I sites was amplified from cDNA library of HEK293 cells. This tag was cloned into the *Age*I site that lies between Dam and LmnB1. The Deadbox domain was synthesized using overlapping oligonucleotides and cloned upstream of Dam. The entire DD-Dam-hCdt1-LmnB1 fragment was then subcloned into the *Eco*RI and *Bst*BI site on Fu_GFP_hLMNA Puro (unpublished).

To generate Fu-BirA-mCherry-m6A, the m6A was synthesized using codon-optimized, overlapping oligonucleotides that span the last 109 amino acids of the restriction endonuclease *Dpn*I and amplified to be flanked by *Bsr*GI and *Bst*BI restriction sites. The amplicon was then cloned downstream of mCherry in the Fu-mCherry-hLMNA-bsr vector (unpublished) using *Bsr*GI and *Bst*BI, generating Fu-mCherry-m6a. Generation one BirA* flanked by *Nhe*I and *Xba*I was amplified from mycBioID-pBABE-puro, a gift from Kyle Roux, and cloned into the *Nhe*I site upstream of mCherry in Fu-mCherry-m6a.

To generate Fu-BirA*Lap2*β*, Lap2*β* was amplified from cDNA material from MEFs and TA cloned into pGEM-t easy (promega). Lap2*β* along with a linker sequence corresponding to the multiple cloning site of pCDNA3.1 was subcloned downstream of BirA* in Fu_BirA*_hLMNA using the *Xho*I and *Bst*BI restriction sites.

### Cell line generation, reagents, antibodies

MEFs were purchased from ATCC (CRL-2752) and cultured according to their establish protocols. These MEFs were transduced with lentiviral particles from the described plasmids. Specifically, viruses were generated in HEK 293T/17 cells (ATCC CRL-11268) by co-transfecting VSV-G, delta 8.9, and the indicated construct. 10 mM sodium butyrate was then added to the transfected cells 3 hours post transfection for an overnight incubation at 37°C, 5% CO2. The transfection media containing sodium butyrate was removed the following day and the cells were washed with 1X PBS. Opti-MEM was then added back to the cells which were then incubated at 37°C, 5% CO_2_. Viral supernatant was collected every 12 hours (up to 4 collections) and the supernatant of all collections were pooled. MEFs were incubated overnight with fresh viral supernatants supplemented with 4 μg/mL polybrene and 10% FBS. Fresh MEF media was then added to the cells after the virus was removed and selected with 10 μg/ml blasticidin or 1μg/ml Puromycin (or both). Antibodies used in this study include: an anti-LMNB antibody (sc-6217, goat IgG; Santa Cruz Biotechnology, Inc.), Alexa Fluor 647 AffiniPure Donkey Anti-Goat (Jackson Immunoresearch, #705-606-147) and anti-biotin antibody (Bethyl Laboratories, Inc. A150-109A).

### Imaging and Immunofluorescence

Cells were prepared for immunofluorescence by plating on sterilized 25-mm round coverslips (German borosilicate glass #1.5; Harvard Apparatus) in 6-well tissue culture dishes. Immunofluorescence was carried out as previously described [12,108]. The nuclear lamina was visualized using an anti-LMNB antibody (sc-6217, goat IgG; Santa Cruz Biotechnology, Inc.) and Alexa Fluor 647 AffiniPure Donkey Anti-Goat (Jackson Immunoresearch, #705-606-147) for secondary detection. Biotinylation was detected using streptavidin-488 (#016-540-084, Alexa Fluor 488 Streptavidin, Jackson Immunoresearch). Immunofluorescence samples were mounted in SlowFade gold (Life Technologies). All Imaging was performed on an inverted fluorescence microscope (AxioVision; Carl Zeiss) fitted with an ApoTome and camera (AxioCam MRm; Carl Zeiss). The objective lens used was a 63× Apochromat oil immersion (Carl Zeiss) with an NA of 1.5 (Immersol 518; Carl Zeiss). All immunofluorescence was performed at room temperature on #1.5 coverslips.AxioVision software (Carl Zeiss) was used for image acquisition. Images were exported as TIFFs to (FIJI ImageJ,National Institutes of Health) for further analyses. [109]

### BioID with BioSITe

NIH3T3 cells expressing LAP2β-BioID, BirA*-m6A-tracer alone, BirA*-m6A-tracer + DD-Dam-LMNB1 constructs were cultured overnight with 50 *μ*M exogenous biotin. Cells expressing BirA*-m6A-tracer + DD-Dam-LMNB1 were cultured with 2 uM shield-1 ligand (AOBIOUS, #AOB1848) for 24 hours prior to the addition of exogenous biotin for a total of 48 hours. Cells were trypsinized, washed in large volume PBS washes, then resuspended in a hypotonic lysis buffer (5 mM PIPES, 85 mM KCL, 1% NP-40, protease inhibitors) for 10 minutes to separate cytoplasmic fraction from nuclear fraction. The resulting nuclei were pelleted and protein extraction was carried out by sonication (three rounds, duty cycle 30%, 20 s pulses) in 50 mM TEABC and 8 M urea. The protein concentration of samples was measured by BCA assay. A total of 10 mg of lysate per replicate was then reduced and alkylated by incubation with 10 mM DTT for 30 min followed by 20 mM IAA for 30 min in the dark. The lysate was diluted to 2 M urea by adding three cell lysate volumes of 50 mM TEABC. The proteins were digested with trypsin (1:20 of trypsin to protein) at 37 °C overnight. The resulting tryptic peptides were desalted using a Sep-PAK C_18_ column and subsequently lyophilized. Protein G agarose beads (Millipore Sigma, #16-266) were washed twice with PBS and 100 *μ*g of anti-biotin antibody (Bethyl Laboratories, Inc. A150-109A) were coupled to 120 *μ*L of protein G bead slurry, overnight at 4 °C. Antibody-coupled beads were further washed with PBS once and BioSITe capture buffer (50 mM Tris, 150 mM NaCl, 0.5% Triton X-100) twice. Lyophilized peptides were dissolved in 1 mL of BioSITe capture buffer and pH solution was adjusted to neutral (7.0 to 7.5). Peptides were subsequently incubated with anti-biotin antibody-bound protein G beads for 1 h at 4 °C. The bead slurry was washed three times with BioSITe capture buffer, three times with 50 mL of Tris, and two times with ultrapure water. Biotinylated peptides were eluted with four rounds of 200 *μ*L elution buffer (80% acetonitrile and 0.2% trifluoroacetic acid in water). The eluents were dried, desalted, and concentrated using homemade C_18_ reversed-phase column.

### Mass Spectrometry

The fractionated peptides were analyzed on an Orbitrap Fusion Lumos Tribrid Mass spectrometer coupled to the Easy-nLC 1200 nanoflow liquid-chromatography system (Thermo Fisher Scientific). The peptides from each fraction were reconstituted in 20 *μ*L of 0.1% formic acid and loaded onto an Acclaim PepMap 100 Nano-Trap Column (100 *μ*m × 2 cm, Thermo Fisher Scientific) packed with 5 *μ*m C18 particles at a flow rate of 4 *μ*L per minute. Peptides were separated at 300 nL/min flow rate using a linear gradient of 7 to 30% solvent B (0.1% formic acid in 95% acetonitrile) over 95 min on an EASY-Spray column (50 cm × 75 *μ*m ID, Thermo Fisher Scientific) packed with 2 *μ*m C18 particles, which was fitted with an EASY-Spray ion source that was operated at a voltage of 2.3 kV.

Mass-spectrometry analysis was carried out in a data-dependent manner with a full scan in the mass-to-charge ratio (*m*/*z*) range of 300–18000 in the “Top Speed” setting, three seconds per cycle. MS and MS/MS were acquired for the precursor ion detection and peptide fragmentation ion detection, respectively. MS scans were measured at a resolution of 120,000 (at *m/z* of 200). MS/MS scans were acquired by fragmenting precursor ions using the higher energy collisional dissociation (HCD) method and detected at a mass resolution of 30,000 (at *m/z* of 200). Automatic gain control for MS was set to one million ions and for MS/MS was set to 0.05 million ions. A maximum ion injection time was set to 50 ms for MS and 100 ms for MS/MS. MS data were acquired in profile mode and MS/MS data in centroid mode. Higher energy collisional dissociation was set to 32 for MS/MS. Dynamic exclusion was set to 35 seconds, and singly charged ions were rejected. Internal calibration was carried out using the lock mass option (*m/z* 445.1200025) from ambient air.

### Database searching, quantification, post processing of MS data

Proteome Discoverer (v 2.2; Thermo Scientific) suite was used for quantitation and identification of peptides from LC–MS/MS runs. Spectrum selector was used to import spectrum from raw file. During MS/MS preprocessing, the top 10 peaks in each window of 100 *m/z* were selected for database search. The tandem mass spectrometry data were then searched using SEQUEST algorithm against protein databases (mouse NCBI RefSeq 73 (58039 entries) with the addition of fasta file entries for BirA*-m6A-tracer and LAP2β-BioID constructs) with common contaminant proteins. The search parameters for identification of biotinylated peptides were as follows: (a) trypsin as a proteolytic enzyme (with up to three missed cleavages); (b) peptide mass error tolerance of 10 ppm; (c) fragment mass error tolerance of 0.02 Da; and (d) carbamido-methylation of cysteine (+57.02146 Da) as a fixed modification and oxidation of methionine (+15.99492 Da) and biotinylation of lysine (+226.07759 Da) as variable modifications. Peptides and proteins were filtered at a 1% false-discovery rate (FDR) at the PSM level using percolator node and at the protein level using protein FDR validator node, respectively. For the MS1 level quantification of the peptides the Minora Feature Detector, using the program’s standard parameters, was used and all of the raw files from the two replicates were quantified together. Unique and razor peptides both were used for peptide quantification, while protein groups were considered for peptide uniqueness. Identified protein and peptide spectral match (PSM) level data were exported as tabular files from Proteome Discoverer 2.2. We used an in-house Python script to compile the peptide level site information mapped to RefSeq databases. We eliminated all non-biotinylated peptides from our analysis. The summary count on the number of supported peptides, PSMs, number of biotinylation sites and quantification was then calculated at the protein level. Quantitation of replicate agreement was plotted and average values between replicates were calculated for total biotinylated protein abundance. Transmembrane domain analysis was carried out as previously described (PMID:28156110). We obtained annotated transmembrane domains, topological domains and subcellular localization from Uniprot. Sites of biotinylation were then compared to annotated topologies to identified the location of biotinylation with respect to lumun, nucleoplasmic and cytoplasmic portions of proteins. For gene set enrichment analysis gene lists were uploaded to the web portal Enrichr (http://amp.pharm.mssm.edu/Enrichr/)(PMID: 27141961).

### DamID

DamID was performed as described previously [9,10,12,20,69]. Cells were transduced with murine retroviruses harboring the Dam constructs. Self-inactivating retroviral constructs pSMGV Dam-V5 (Dam-Only), pSMGV Dam-V5-Lamin A (Dam-Lamin A), pSMGV Dam-V5-Lap2*β* (Dam-Lap2*β*) and pSMGV Dam-V5-EMD (Dam-EMD) were transfected using Fugene 6 transfection reagent (Promega, E2691) into the Platinum-E packaging line (Cell Biolabs, RV-101) to generate infectious particles. These viral supernatants in DMEM complete media were used to directly infect MEF lines. Retroviral infections were carried out by incubating MEFs overnight with either Dam-only, Dam-LmnB1, Dam-Lap2*β* or Dam-EMD viral supernatant and 4 μg/mL polybrene. Cells were allowed to expand for 2-4 days then pelleted for harvest.

MEFs were collected by trypsinization and DNA was isolated using QIAamp DNA Mini kit (Qiagen, 51304), followed by ethanol precipitation and resuspension to 1 μg/ul in 10 mM Tris, pH 8.0. Digestion was performed overnight using 0.5-2.5 μg of this genomic DNA and restriction enzyme DpnI (NEB, R0176) and then heat-killed for 20 minutes at 80°C. Samples were cooled, then double stranded adapters of annealed oligonucleotides (IDT, HPLC purified) AdRt (5′ -CTAATACGACTCACTATAGGGCA GCGTGGTCGCGGCCGAGGA-3′) and AdRb (5′ -TCCTCGGCCG-3′) were ligated to the DpnI digested fragments in an overnight reaction at 16°C using T4 DNA ligase (Roche, 799009). After incubation the ligase was heat-inactivated at 65°C for 10 minutes, samples were cooled and then digested with DpnII for one hour at 37°C (NEB, R0543). These ligated pools were then amplified using AdR_PCR oligonucleotides as primer (5′ -GGTCGCGGCCGAGGATC-3′) (IDT) and Advantage cDNA polymerase mix (Clontech, 639105). Amplicons were electrophoresed in 1% agarose gel to check for amplification and the size distribution of the library and then column purified (Qiagen, 28104). Once purified, material was checked for LAD enrichment via qPCR (Applied Biosystems, 4368577 and StepOne Plus machine) using controls specific to an internal Immunoglobulin heavy chain (Igh) LAD region (J558 1, 5′-AGTGCAGGGCTCACAGAAAA-3′, and J558 12, 5′-CAGCTCCATCCCATGGT TAGA-3′) for validation prior to sequencing.

### DamID-seq Library Preparation and Sequencing

In order to ensure sequencing of all DamID fragments, post-DamID amplified material was randomized by performing an end repair reaction, followed by ligation and sonication. Specifically, 0.5-5 μg of column purified DamID material (from above) was end-repaired using the NEBNext End Repair Module (NEB E6050S) following manufacturer’s recommendations. After purification using the QIAquick PCR Purification Kit (Qiagen, 28104), 1μg of this material was then ligated in a volume of 20 μL with 1μl of T4 DNA ligase (Roche, 10799009001) at 16°C to generate a randomized library of large fragments. These large fragments were sonicated (in a volume of 200μL, 10mM Tris, pH 8.0) to generate fragments suitable for sequencing using a Bioruptor^®^ UCD-200 at high power, 30 seconds ON, 30 seconds OFF for 1 hour in a 1.5 mL DNA LoBind microfuge tube (Eppendorf, 022431005). The DNA was then transferred to 1.5 ml TPX tubes (Diagenode, C30010010-1000) and sonicated for 4 rounds of 10 minutes (high power, 30 seconds ON and 30 seconds OFF). The DNA was transferred to new TPX tubes after each round to prevent etching of the TPX plastic. The sonication procedure yielded DNA sizes ranging from 100-200 bp. After sonication, the DNA was precipitated by adding 20 μl of 3M sodium acetate pH 5.5, 500 μl ethanol and supplemented with 3 μl of glycogen (molecular biology grade, 20 mg/ml) and kept at −80°C for at least 2 hours. The DNA mix was centrifuged at full speed for 10 min to pellet the sheared DNA with the carrier glycogen. The pellet was washed with 70% ethanol and then centrifuged again at full speed. The DNA pellet was then left to air dry. 20 μl of 10 mM Tris-HCl was used to resuspend the DNA pellet. 1 μl was quantified using the Quant-iT PicoGreen dsDNA kit (Invitrogen, P7589). Sequencing library preparation was performed using the NEBNext Ultra DNA library prep kit for Illumina (NEB, E7370S), following the manufacturer’s instructions. Library quality and size was determined using a Bioanalyzer 2100 with DNA High Sensitivity reagents (Agilent, 5067-4626). Libraries were then quantified using the Kapa quantification Complete kit for Illumina (Kapa Biosystems, KK4824) on an Applied Biosystems 7500 Real Time qPCR system. Samples were normalized and pooled for multiplex sequencing.

### DamID-seq Data Processing

DamID-seq analysis—DamID-seq reads were processed using LADetector (available at https://github.com/thereddylab/LADetector), an updated implementation of LADetector described in Harr et al. (2015) with incorporated sequence mapping. Specifically, 5’ ends of reads were quality trimmed using a sliding window quality score average over 3 bases and a minimum score cutoff of 30. This was followed by trimming any matching overlap between read-ends and sequencing or DamID adaptor-primer sequence. Reads containing a DamID adaptor-primer sequence were split and adaptor-primer sequence removed. All resulting reads greater than 20 bp were aligned to mm9 using Bowtie (Langmead et al., 2009) with parameters “—tryhard–best–m 1.” Unaligned reads had 13 bases trimmed from the 5′ end and remapped, and the resultant unmapped reads were trimmed 13 bases from the 3′ end and remapped. Total aligned reads were assigned to *Dpn*I bins, with reads straddling bin boundaries counting toward both. Prior to scoring, a value of 0.5 was added to bins with no reads. Bins falling in unaligned regions were removed prior to analysis. DamID scores were calculated for all non-zero bins as the log2 ratio of Dam-Lamin B1 over unfused Dam. Scores were partitioned using circular binary segmentation using the DNAcopy package in R (Seshan and Olshen, 2018). LADs were classified as regions > 100 kb in size of positive signal, allowing for smaller regions of negative signal < 10 kb in size.

## Data Availability

The mass spectrometry proteomics data have been deposited to the ProteomeXchange Consortium (http://proteomecentral.proteomexchange.org) via the PRIDE partner repository with the data set identifier PXD012943.

The sequencing data have been deposited in Gene Expression Omnibus (GEO): GSE128239

## Supplemental Material

**Figure S1:**
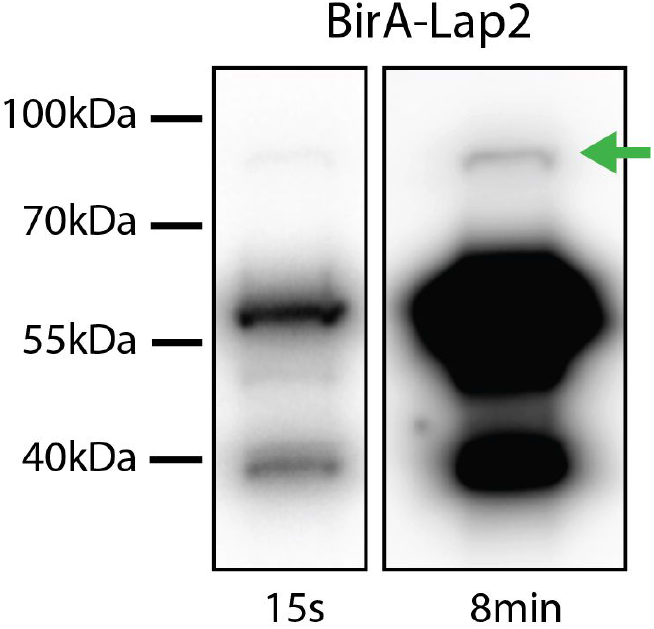
Western blot showing relative levels of BirA-Lap2β levels at 2 different exposures (15 seconds, 8minutes). BirA-Lap2β fusion protein is indicated by the green arrow and is expressed at 1-2% of endogenous Lap2β. (Antibody BD611000)

**Figure S2:**
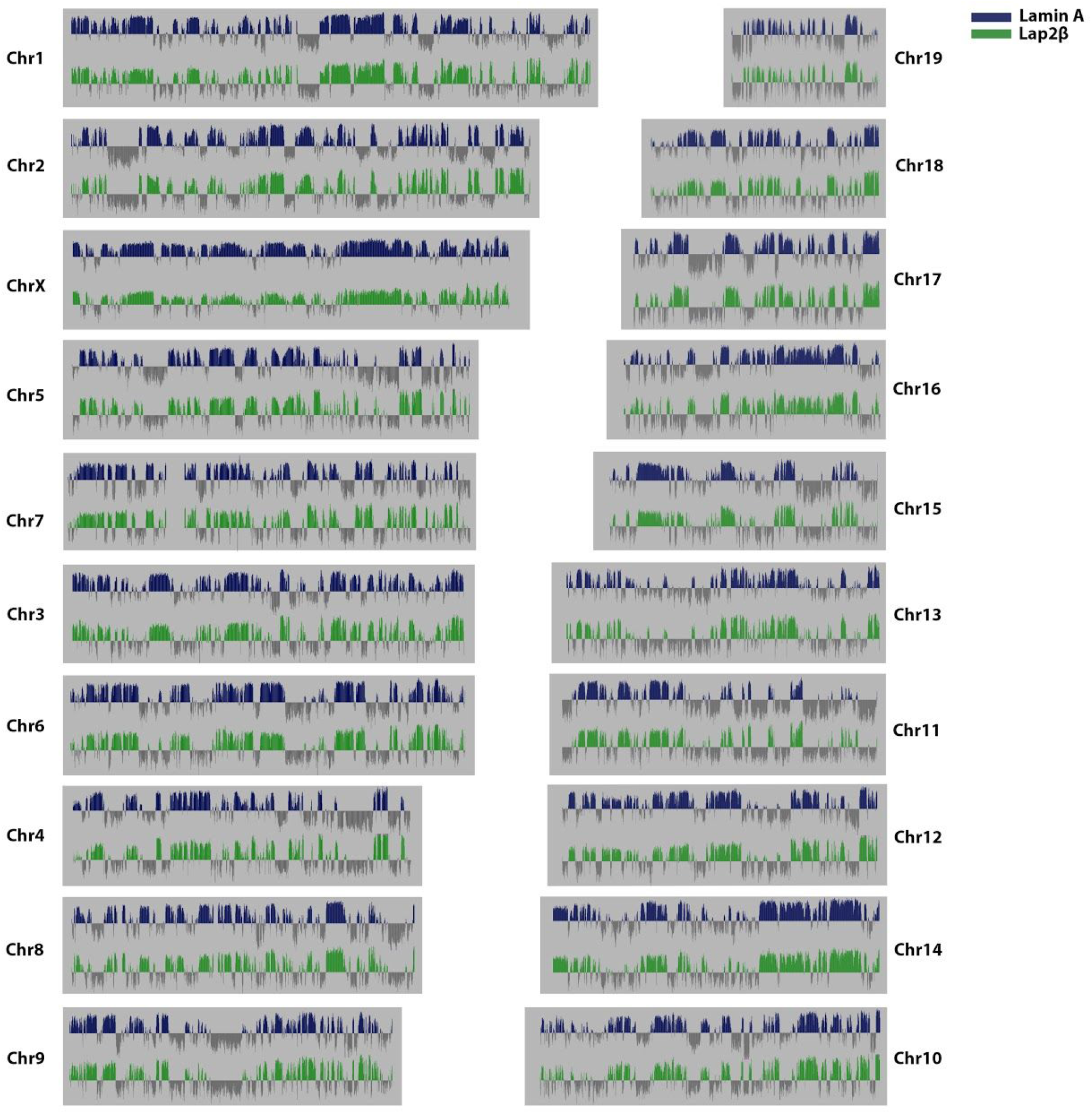
Genome wide Lamin A (blue) and Lap2β (green) DamID log2 ratio plots.

